# Ensemble Coding of Hidden Objects in Visual Cortex

**DOI:** 10.64898/2025.12.29.696959

**Authors:** Shude Zhu, Danielle A. Lopes, Stephen N. Cital, Tirin Moore

## Abstract

Many species exhibit the understanding that visual objects that become hidden by others nonetheless still exist, a property known as object permanence. Previous studies in human and nonhuman primates have provided evidence that neurons within visual cortex encode objects that are remembered but not seen. However, past neurophysiological studies have generally failed to find evidence of visual cortical representations of hidden objects. We measured the activity of large populations of neurons within dorsal extrastriate cortex of macaques trained to monitor the identity of visual objects that moved behind an irrelevant occluder. We found that although the firing rates of neuronal populations signaled the trajectory of hidden objects throughout the occlusion period, coding of object identity in the same activity decayed to chance before the behavioral trial ended. Nevertheless, information about the hidden object was present in the coordinated activity of neuronal populations. Specifically, the strength and presence of pairwise cross-correlations reliably depended on the identity of the hidden object. These results demonstrate that ensembles of visual cortical neurons preserve information about hidden objects independent of single neuron firing rates.

## Introduction

Important visual objects are often temporarily hidden from view. Nonetheless, many species of birds and mammals demonstrate the understanding that objects persist despite discontinuities in their visibility (de Bois & Novak, 1994; Schaffer et al., 2024), a capacity known as object permanence (Piaget, 1954). Although it is clear that robust representations of hidden objects must and do exist in the brain, e.g. prefrontal cortex (Fuster & Alexander, 1971), the extent to which sensory networks contribute to those representations remains a mystery. Numerous studies have demonstrated the persistence of neural representations of unseen visual stimuli, whether imagined (Pearson, 2019; Sakai & Miyashita, 1991; Schlack & Albright, 2007) or remembered (Christophel et al., 2017). In the latter case, studies in both human and nonhuman primates have yielded evidence that neurons within visual cortex encode information about behaviorally relevant objects that are remembered but not seen (Harrison & Tong, 2009; Miller et al., 1993; Miyashita & Chang, 1988; Serences, 2016; Serences et al., 2009). For example, neurophysiological studies in nonhuman primates demonstrate that neurons at early to middle stages of visual cortex not only signal the locations of targets held in working memory (Jonikaitis et al., 2025; O’Herron & von der Heydt, 2009; Super et al., 2001; van Kerkoerle et al., 2017), but also the featural properties of those targets (Bisley et al., 2004; Dotson et al., 2018; Huang et al., 2024; Mendoza-Halliday et al., 2014; Motter, 1994; Panichello & Buschman, 2021; Yiling et al., 2024; Zaksas & Pasternak, 2006). This evidence establishes that the coding of visual objects can be achieved, at least to some extent, by visual cortical neurons even in the absence of retinal stimulation.

The question of the visual system’s involvement in representing hidden objects is more complex, however. In the absence of competing input, neuronal responses to recently seen, behaviorally relevant stimuli can persist in visual cortex, presumably due to top-down inputs from other structures (Bichot et al., 2015; Bichot et al., 2019). Once occluded, however, sensory input from the occluder should dominate the output of visual cortical neurons. Indeed, it has been shown that representations of objects during working memory in prefrontal cortex persist in the presence of intervening stimuli, whereas they are disrupted in temporal visual cortex (Miller et al., 1996; Woloszyn & Sheinberg, 2009). This could mean that visual cortical neurons generally do not robustly encode information about hidden objects. On the other hand, neurophysiological studies within dorsal visual areas, where neurons are more specialized for motion and spatial processing (Born & Bradley, 2005; Kravitz et al., 2011), have revealed compelling evidence that many neurons signal the presence of occluded objects. For example, neurons within posterior parietal cortex signal the motion of targets behind an occluder (Assad & Maunsell, 1995), and neurons within superior temporal cortex appear selective to the presence or absence of occluded stimuli (Baker et al., 2001). Such evidence raises the possibility that visual cortical areas more involved in tracking the position of objects might in some way also contribute to representing the identity of tracked objects. To address this possibility, we recorded the activity of neurons within the superior temporal sulcus (STS) of monkeys trained to track visual objects that were briefly hidden behind an occluder. In each experiment, the activity of large neuronal populations was measured simultaneously thus allowing us to assess the extent to which those populations could collectively signal information about the occluded object.

## Results

We recorded the activity of large populations of neurons within the STS of macaque monkeys trained to track the identity of moving visual objects (METHODS). Two monkeys (Hd, 17 kg; Hb, 12 kg) performed a task in which they were rewarded for detecting changes in visual objects that moved across a display (Figure 1). After the monkey fixated a central spot, a single visual object appeared at a randomly chosen location. Shortly thereafter, the object moved along a trajectory that intersected a fixed location. On some trials, the object was replaced with another, and the monkey received a juice reward if it shifted its gaze to the new object. On other trials, the object did not change, and the monkey was rewarded for maintaining fixation. On 20-30% of the total trials, the object was visible throughout its trajectory. On most trials however, the object was briefly hidden when it moved behind a static, opaque occluder (Figure 1a). On these trials, changes in the object always occurred behind the occluder. In one version of the task (monkey Hd), the moving object stopped behind the occluder (1 s) before it was revealed when the occluder disappeared. In the other version of the task (monkey Hb), the moving object kept moving behind the occluder for a period (∼0.7 s) until it emerged. In either case, the monkey was rewarded for remembering the hidden object. In a single session, four to six different objects moved along two to four different trajectories that intersected the fixed occluder location (Figure 1b). Both monkeys reliably detected identity changes on occlusion trials (d’=1.2, Hd; d’=0.7, Hb) (Figure 1c) and were nearly perfect on trials when the object remained visible (d’=3.9, Hd; d’=3.2, Hb) (METHODS).

**Figure 1.**
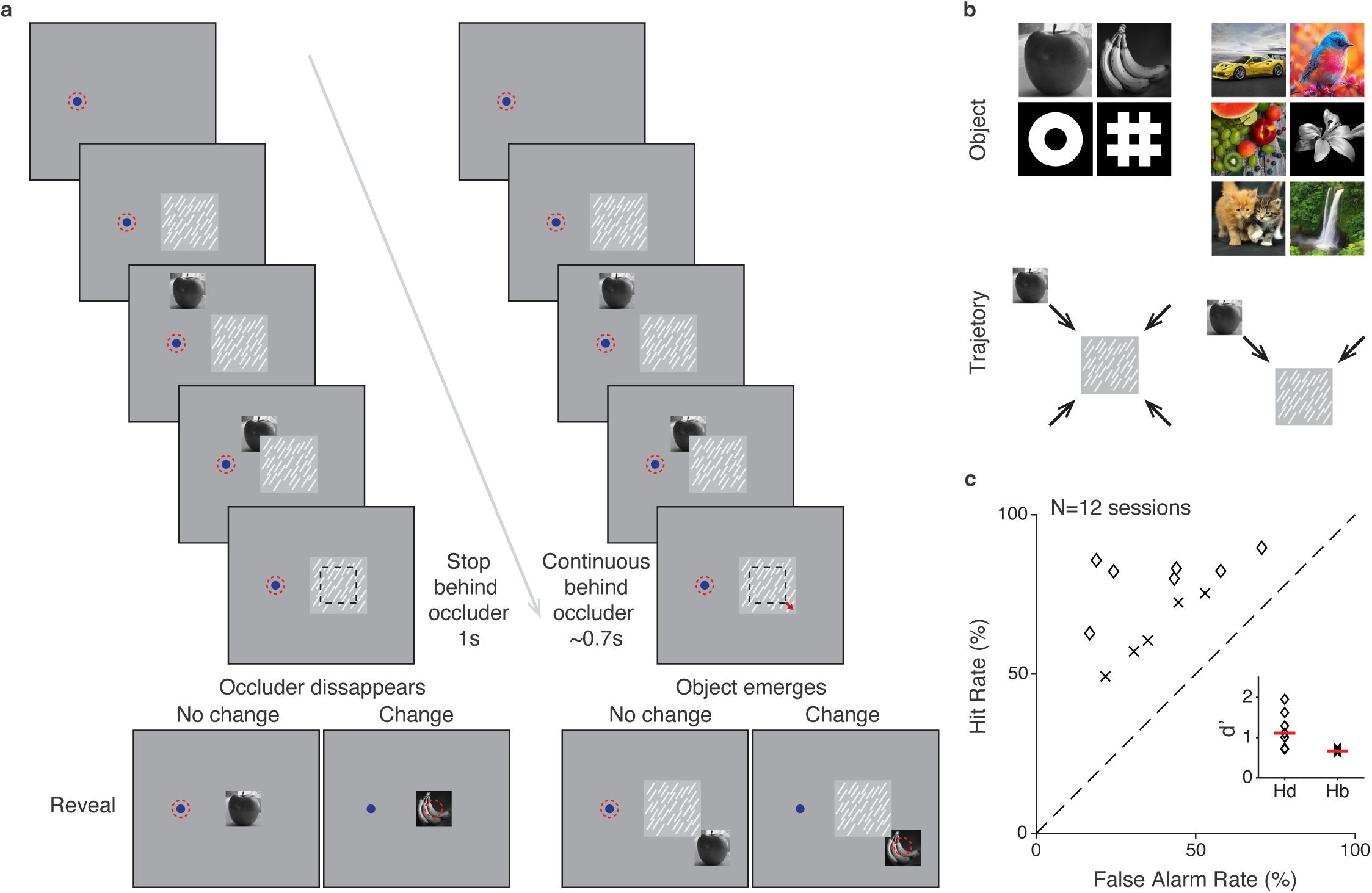
Tracking changes to briefly hidden objects. **a**, Sequence of task events. After central fixation, a static occluder (textured square) appeared, followed by the appearance of a target object (apple). The object then moved toward the occluder and either stopped for a period until the occluder disappeared (Task 1) or continued moving until reemerging from behind the occluder (Task 2). While occluded, the object could change identity. After the object was revealed, monkeys were rewarded for shifting its gaze to the object on change trials, or for maintaining fixation on no-change trials. **b**, Example stimulus sets and trajectories. Across sessions, four different target (grayscale) objects moved along four trajectories (Task 1) or six different (colored) objects moved along two trajectories (Task 2). **c**, Behavioral performance of the two monkeys at detecting changes to the briefly hidden objects: hit rates, false alarm rates and sensitivity (d’). Each symbol denoted data from each session; horizontal lines denote mean d’.

During the task, the activity of single neurons within the STS, particularly areas MT and MST, was recorded using Neuropixels probes (METHODS) (Figure 2a). In each session, neuronal receptive fields (RFs) were distributed across locations within the contralateral visual field due to the nonorthogonal angle of probe penetration into the cortex. That distribution, which comprises the population RF was used to determine the location of the occluder (Figure 2b). On occlusion trials, neurons responded to the occluder, to the appearance and movement of the target object, and to the object’s reappearance after occlusion; responses were heterogeneous, consistent with variation in RF position and properties (Figure 2c). In the example session (211 neurons), mean population firing during the period of full occlusion differed across trajectories (Friedman test, *P* < 0.001), but not across the four object identities (Friedman test, *P* = 0.21) (Figure 2d).

**Figure 2.**
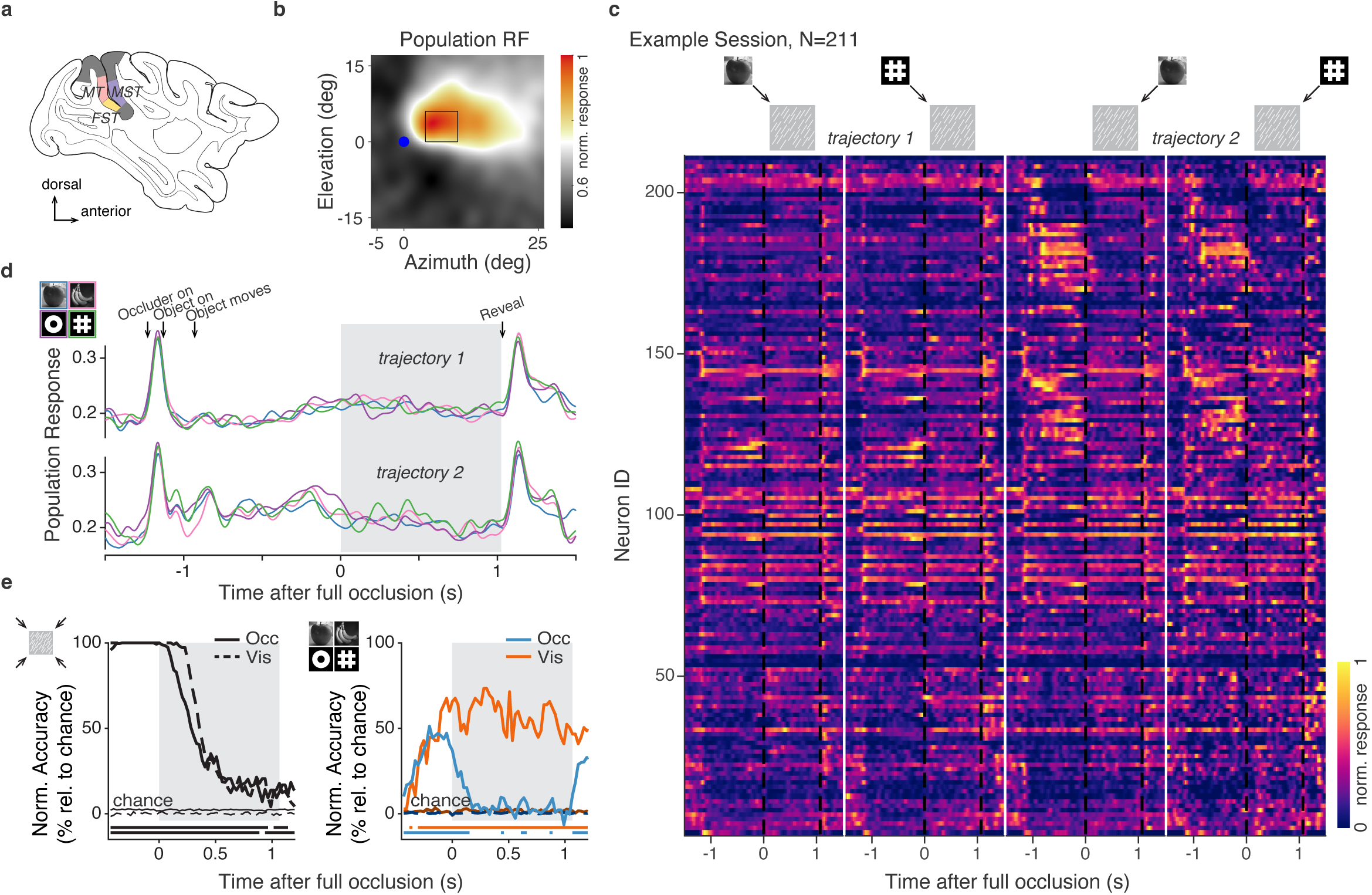
STS population activity during tracking of briefly hidden objects. **a**, Estimated recording locations in STS shown on a sagittal section; areas MT, MST, and FST are indicated. **b**, Population RF map from an example recording session showing mean normalized response of 211 single neurons in STS. Open square shows the location of the occluder; blue dot denotes the fixation point. **c**, Trial-averaged, normalized responses of individual neurons (row) for two objects by two trajectory conditions in Task 1. Vertical dashed lines mark the start and end of full occlusion. **d**, Mean normalized population response for the example session, separated by trajectory (rows) and object identity (colors). Shading indicates the period of full occlusion during the occluded conditions. **e**, Time-resolved decoding of trajectory (left) and object identity (right) from population activity for occlusion trials (solid) during the occlusion period and visible trials (dashed) during corresponding interval. Performance is plotted as normalized accuracy relative to theoretical chance: (acc-chance)/(1-chance) *100. Thin traces show shuffled-label controls. Thick horizontal lines along x-axis denote time bins with decoding significantly above shuffled controls (shuffle test; METHODS).

To better quantify population sensitivity to hidden trajectories and object identities, we trained time-resolved logistic regression classifiers on the population activity (METHODS). In the example session, decoding of the four trajectories was nearly perfect, but decayed rapidly after occlusion onset (Figure 2e). Nevertheless, decoding remained above chance throughout the occlusion interval until object was revealed (Exponential fit; final-bin accuracy above chance: 11.8%, 95% CI 9.8−13.8%; asymptote: 10.2%, 95% CI 7.5−12.8%; *t_95_* = 822 ms, 95% CI 740−904 ms, where *t_95_* denotes the time to reach 95% of the decay from peak to asymptote). A comparable persistence was observed in visible condition in which the object entered the population RF and then stopped for the rest of the trial (final-bin accuracy above chance: 14.0%, 95% CI 11.9−16.0%; asymptote: 13.8%, 95% CI 11.7−16.0%; *t_95_* = 631 ms, 95% CI: 580−682 ms). In both cases, population activity continued to distinguish trajectories through trial end.

In contrast, the same population failed to signal the identity of the hidden objects. In the visible condition, identity decoding rose as the object entered the population RF and plateaued at ∼55%, confirming sufficient object selectivity in the population. However, when the object became hidden behind the occluder, identity decoding declined rapidly to near chance and remained there until reveal (asymptote: 3.0%, 95% CI 1.6−4.5%; *t_95_* = 278 ms, 95% CI 183−372 ms). At asymptote, decoding was not consistently above shuffled controls across time (shuffle test, 100 permutations; *P* > 0.05 for the majority of time bins). Thus, in this session, STS population activity robustly signaled the hidden trajectory but showed little evidence for sustained coding of hidden object identity.

Across 12 sessions in two monkeys, we measured the activity of 1,518 neurons within the STS (78-211 neurons per session). For each session, we quantified population sensitivity to hidden trajectories using time-resolved classifiers trained to decode the trajectory condition (Figure 3a). As in the example session, trajectory decoding declined at full occlusion onset but remained above chance throughout the occlusion interval. We observed this in both task variants, whether the object remained behind the occluder (Task 1) or continued moving and subsequently re-emerged. In the visible conditions, trajectory decoding remained high throughout the trial in Task 2 as the object remained in motion. In spite of the task differences, the time course of trajectory decoding during occlusion was similar up to the time of reveal/emergence. In both tasks, decoding decayed but remained above chance through the end of the occlusion period (Task 1: final-bin accuracy above chance: 11.9%, 95% CI 10.2−13.5%; asymptote: 9.7%, 95% CI 7.3−12.0%; *t_95_* = 881 ms, 95% CI 807−954 ms; Task 2: final-bin accuracy above chance: 17.6%, 95% CI 15.0−20.2%; asymptote: 10.0%, 95% CI 3.9−16.0%; *t_95_* = 831 ms, 95% CI 718−945 ms). Performance in the combined activity during the late occlusion period was similar in the two tasks and across recording sessions (Task 1: 33.3% ± 4.8%; Task 2: 35.1% ± 3.5%; mean ± s.e.m. across sessions; *P* < 0.01 in each session) (Figure 3b) (METHODS).

**Figure 3.**
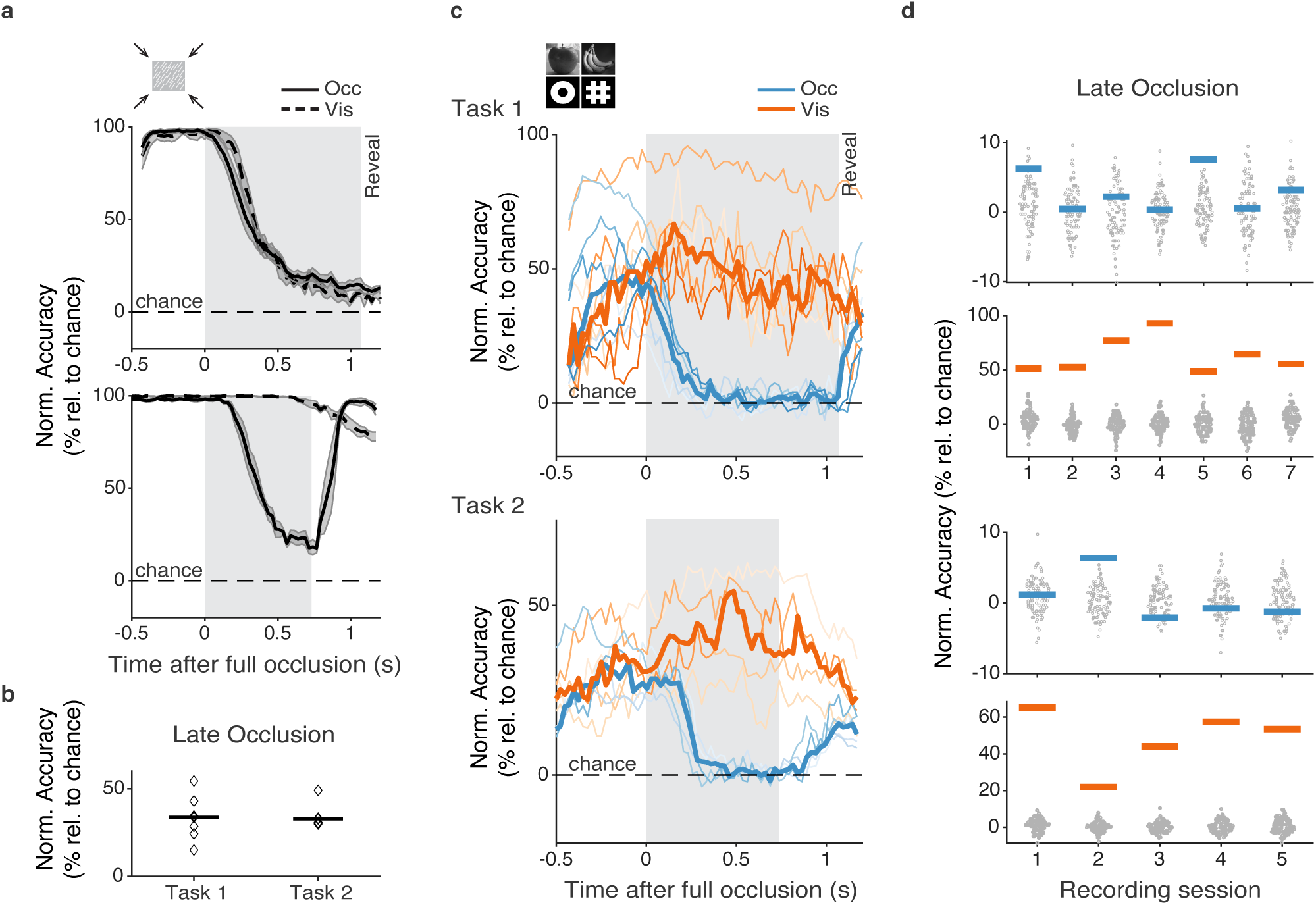
Activity of STS neurons during tracking briefly hidden objects. **a**, Time-resolved decoding of object trajectory across recording sessions for Task 1 (top) and Task 2 (bottom) during visible (dashed) and occlusion (solid) conditions. Shading around performance mean denotes s.e.m. across sessions. **b**, Mean trajectory decoding performance during the late occlusion period of the occlusion conditions. **c**, Time-resolved decoding of object identity across sessions for Task 1 (top) and Task 2 (bottom) during visible and occlusion conditions. **d**, Mean object decoding performance (horizontal lines) for Task 1 (top) and Task 2 (bottom) in each session during the late occlusion period of the occlusion conditions and during the corresponding time window of the visible conditions. Gray points show shuffle distributions (n=100 permutations) for each session. Other conventions as in previous figures.

Having established that populations of STS neurons could signal object trajectories throughout occlusion, we next asked whether information about object identity similarly persisted while the object was hidden. Results from the example session suggested the contrary though they need not be representative of the full dataset. Thus, we trained classifiers in each of the 12 sessions to decode object identity (Figure 3c). In the visible condition, identity decoding rose as the object entered the population RF and then remained relatively stable. In each recording session, activity measured during the late period of the visible conditions could be used to decode object identity (Task 1: 63.3% ± 6.2%; Task 2: 48.4% ± 7.4%; mean ± s.e.m.; *P* < 0.01 in each session).

In contrast to the visible condition, across sessions, we failed to find consistent evidence that neurons continued to signal the identity of objects once they became hidden. As in the example, mean decoding performance decayed to near chance levels within ∼380 ms where it remained until the object was revealed (Task 1: asymptote: 1.0%, 95% CI 0.3−1.7%; *t_95_* = 382 ms, 95% CI 343-420 ms; Task 2: asymptote: 1.0%, 95% CI 0.2−1.9%; *t_95_* = 374 ms, 95% CI 334-413 ms). In most recording sessions (10/12), in contrast to the visible conditions, decoding of object identity did not exceed chance performance during the late period of occlusion (*P* > 0.05) (Figure 3d). This was true in both versions of the task. Only two sessions (one per task) showed weak but significant above-chance identity decoding during the late occlusion period (Task 1, session 5, *P* < 0.01; Task 2, session 2, *P* < 0.01). Thus, across sessions, populations of STS neurons consistently signaled the trajectory of hidden objects, but not their identity.

The absence of reliable decoding in the late occlusion period suggests that information about the identity of hidden objects is not conveyed by the firing rates of STS neurons. We therefore asked whether identity might instead be expressed in the coordinated activity of neuronal populations on finer temporal scales (<100 ms). That information would likely be obscured by the summation of spike counts into larger decoding bins. Indeed, evidence indicates that the maintenance of information during working memory is achieved both by persistent spiking activity and by short-term plasticity (Lundqvist et al., 2018; Mongillo et al., 2008; Panichello et al., 2024; Stokes, 2015). In the latter case, in the absence of persistent activity, working memory is instead represented by memorandum-specific ensembles of neurons. Evidence of this can be observed in the coordinated activity between neurons on short timescales, such as in the pairwise spike cross-correlations in neuronal activity (Panichello et al., 2024). Thus, we looked for similar evidence of stimulus-specific ensembles for hidden objects during the late occlusion.

As in the decoding analyses, we leveraged the Neuropixels recordings to measure functional connections among the large number of simultaneously recorded neuron pairs (8,559 ± 1,536 pairs per session, mean ± s.e.m.). We examined pairwise spike-train cross-correlations to assess their dependence on object identity. For each session and each hidden object condition, we computed the cross-correlograms (CCGs) for all neuronal pairs using spikes during the late occlusion period, following established procedures (METHODS). Our analysis focused on contrasting the CCGs between any combination of two different objects within a session.

Across recordings, 1.8% ± 0.28% of neuronal pairs exhibited significant CCGs (peak >7 SD_noise_), yielding 1,894 significant CCGs across all sessions. When comparing these significant CCGs between two different hidden objects, we observed differences in the pattern of functional connectivity (Figure 4a). Specifically, pairs of neurons could exhibit significant CCGs with one object, but not the other, resulting in two distinct ensembles. These differences reflected systematic changes in CCG peak amplitude across object identities of each neuronal pair. To summarize object-dependent changes across all significant pairs, we computed the Euclidean distance between vectors of CCG peaks of the ensembles observed for the two different objects (Figure 4b) (METHODS). In the example object comparison (Task 1, session 3, 157 neuronal pairs), the distance between the ensembles differed significantly from the shuffled control (Z=2.84; shuffle test, 1000 permutations; *P* = 0.013). However, the distance between the peak lags of the ensembles was not significant (Z=0.65; shuffle test, 1000 permutations; *P* = 0.26).

**Figure 4.**
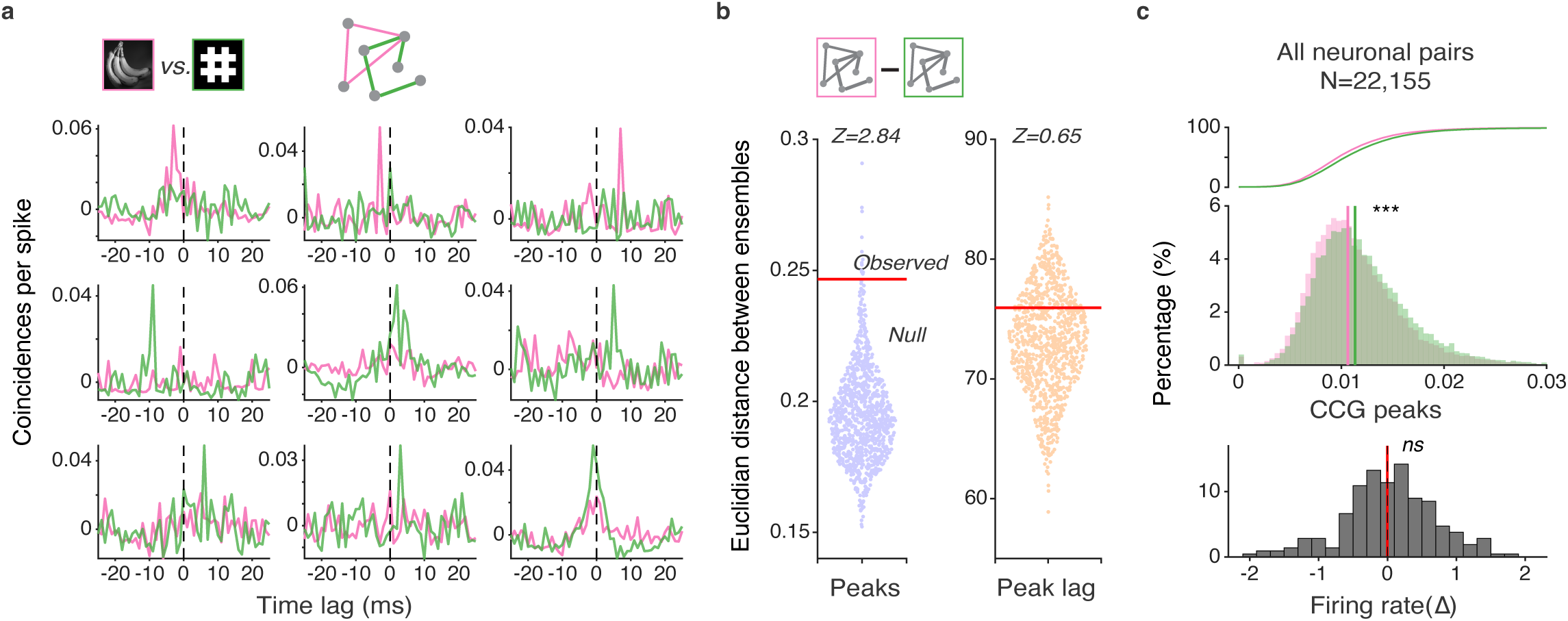
Dependence of neuronal cross-correlations on hidden object identity. **a**, Example spike cross-correlations in the late occlusion period. Shown are the CCGs of a neuronal pair for two different hidden objects (green and pink). In each case, one of the computed CCGs was significant. **b**, Comparison of the Euclidean distance between ensembles for two different hidden objects to the shuffled distribution of distances in an example session. **c**, Comparison of CCG peaks of all neuronal pairs for the two hidden objects (cumulative and count distributions). Bottom plot shows distribution of mean neuronal firing rate differences for the two objects during the late occlusion period. Asterisks denote significance (***, *P* < 0.001; *ns*, nonsignificant).

Next, given the clear differences in CCG peaks observed between objects in significantly correlated neuronal pairs, we asked whether such differences might also be present in the total set of neuronal pairs. For the two objects in the example session, we compared the peaks in the CCGs measured during the late occlusion period using all 22,155 neuronal pairs (Figure 4c). This revealed that the distributions of CCG peaks differed significantly across the two objects and the average CCG peak between neuronal pairs was greater for one object than the other (Wilcoxon signed-rank test*, P* < 0.001). Thus, the likelihood of coincident spikes between pairs of neurons differed reliably between the two hidden objects. Notably, we observed this effect despite an absence of a reliable difference in average firing rate between the two conditions measured in the same epoch (Wilcoxon signed-rank test*, P* = 0.59). This result indicates that the two hidden objects evoked different patterns of coordinated activity between neurons that was independent of their firing rates.

Across recording sessions, we observed a similar pattern of results as the example in both versions of the task and in both monkeys. Specifically, we found differences in the pattern of neuronal cross-correlations between objects when measured either in all neuronal pairs, or pairs with significant CCGs. Across sessions, we first measured the differences in the neuronal ensembles comprised of significantly correlated neuronal pairs (Figure 5a). As in the example, we quantified ensemble separation as the Euclidean distance between vectors of CCG peaks for the two different objects and compared them to the shuffled (null) control (METHODS). Ensemble distances were reliably greater than the null distribution across sessions and tasks (Wilcoxon signed-rank test, *P* < 0.001). Thus, the composition of correlated neuronal ensembles depended reliably on the identity of the hidden object. In addition, across sessions and in both tasks, we also observed object-dependent differences in the peak lags of the ensembles (Wilcoxon signed-rank test; Task 1: *P* < 0.01; Task 2: *P* < 0.001). This latter result suggests that object-specific ensembles may also exhibit different temporal dependences in their correlated spike trains.

**Figure 5.**
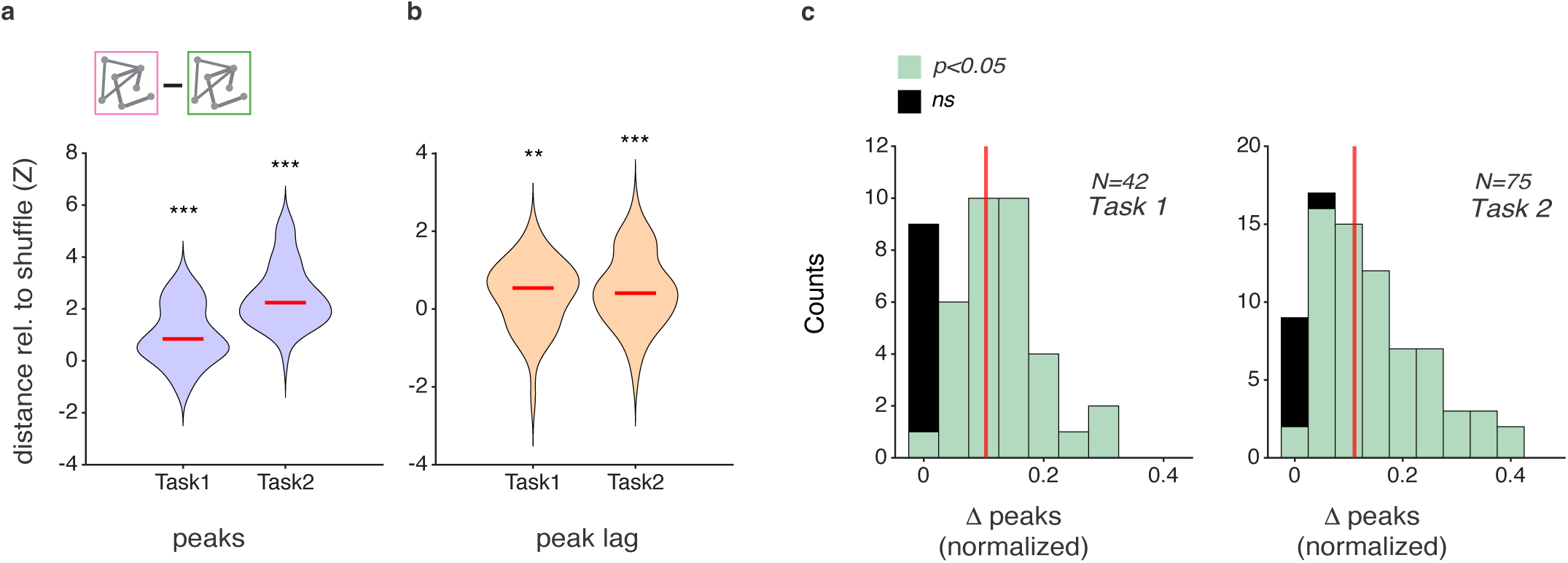
Object-dependent cross-correlations across sessions. **a**, Comparison of the Euclidean distances between ensembles for all sets of two hidden objects to their shuffled distributions across sessions. Mean (red line) and variation (purple) in ensemble peak distances are shown relative to their shuffled control (Z) for both tasks. **b**, Same as **a**, but distances between ensembles is measured in peak lag. **c**, Comparisons of CCG peaks for all sets of two hidden objects (117 total) across sessions and tasks, using the CCGs from all neuronal pairs. Distributions of normalized differences CCG peaks between any two objects are shown for both tasks. Colored elements denote individually significant comparisons; black elements denote nonsignificant ones. Asterisks denote significance (**, *P* < 0.01; ***, *P* < 0.001; *ns*, nonsignificant).

Finally, as in the example session, we looked for differences in correlated activity in the total set of neuronal pairs across all sessions and in both tasks. In each session, we compared the CCG peaks between each combination of two different objects (6 comparisons in Task 1, 15 comparisons in Task 2). This yielded a total of 117 object comparisons across all sessions and tasks. When comparing the CCGs of all neuronal pairs, 101/117 (86%) hidden object comparisons exhibited significant differences in the distribution of CCG peaks (Task 1: 34/42, 81%; Task 2: 67/75, 89%) (Figure 5b) (METHODS). Thus, regardless of the extent to which significant functional connectivity was measurable between neuronal pairs, differences in correlated activity were observable between hidden objects. Specifically, the distribution of CCG peaks, and thus the likelihood of coincident spikes between neuronal pairs, differed reliably between hidden objects in a vast majority of comparisons. Overall, both results demonstrate that despite the absence of reliable differences in average firing rates measured in the late occlusion period, hidden objects nonetheless evoked reliable differences in coordinated activity among populations of STS neurons.

## Discussion

We measured the activity of large populations of STS neurons of macaques trained to monitor the identity of visual objects that moved behind an occluder. Across recording sessions, we found that although populations of STS neurons consistently signaled the trajectory of hidden objects, they could not reliably signal object identity. Given the absence of a clear object signal, we assessed the extent to which coordinated neuronal activity depended on the identity of the hidden object. We leveraged the large numbers of simultaneously recorded neurons to measure cross-correlations among neuronal pairs. We found that information about the hidden object was present in the variation in cross-correlations among neuronal populations. This was evident when comparing the similarity of significantly connected ensembles identified with different objects; those ensembles reliably differed between different hidden objects. We also observed the object dependence in the overall differences in the pairwise cross-correlations among neurons within the entire population. Regardless of the extent to which significant functional connectivity was measurable between neuronal pairs, differences in correlated activity were nonetheless observable between different hidden objects. Our results demonstrate that despite the absence of reliable differences in average firing rates measured in the late occlusion period, hidden objects nonetheless evoked reliable differences in coordinated activity among populations of STS neurons.

Like many other species, macaque monkeys demonstrate the understanding that objects persist despite discontinuities in their visibility (de Bois & Novak, 1994). Indeed, this fact is also exemplified by our behavioral results. However, past neurophysiological studies have generally failed to find evidence of visual cortical representations of hidden objects. At early to middle stages of visual cortex, neurons signal the locations of targets held in working memory (Jonikaitis et al., 2025; O’Herron & von der Heydt, 2009; Super et al., 2001; van Kerkoerle et al., 2017), as well as the featural properties of those targets (Bisley et al., 2004; Dotson et al., 2018; Huang et al., 2024; Mendoza-Halliday et al., 2014; Motter, 1994; Panichello & Buschman, 2021; Yiling et al., 2024; Zaksas & Pasternak, 2006). This evidence establishes that the coding of visual objects can be achieved, at least to some extent, by visual cortical neurons even in the absence of retinal stimulation. Yet, in the presence of other stimuli, such coding is disrupted (Miller et al., 1996; Woloszyn & Sheinberg, 2009). Thus, it has remained unclear to what extent visual cortical neurons can contribute to the representation of occluded stimuli, at least in their individual firing rates. Recent studies of working memory mechanisms suggest that maintenance of information can be achieved by short-term plasticity (Lundqvist et al., 2018; Mongillo et al., 2008; Panichello et al., 2024; Stokes, 2015), and the resulting memorandum-specific neuronal ensembles. Evidence of this can be observed in the coordinated activity between neurons on short timescales, such as in the pairwise spike cross-correlations in neuronal activity (Panichello et al., 2024). Our results suggest that such a mechanism may also exist within cortex and may contribute to the maintenance of visual information for target objects tracked behind occluders.

## Methods

### Experimental model and subject details

Two adult male rhesus macaques (*Macaca mulatta*, Hd, 13 years, 17 kg; Hb, 15 years, 12 kg), participated in the study. All experimental procedures conformed to the National Institutes of Health Guide for the Care and Use of Laboratory Animals, the Society for Neuroscience Guidelines and Policies, and were approved by the Stanford University Institutional Animal Care and Use Committee (IACUC; protocol #APLAC-9900).

### Behavioral task

Experiments were controlled by a Dell Precision Tower 3620 workstation and implemented in custom MATLAB scripts (R2018a; MathWorks) using Psychophysics Toolbox (PTB-3) (Kleiner et al., 2007) and Eyelink toolboxes (Cornelissen et al., 2002). Eye position was monitored online with an EyeLink 1000 chair-mounted system (SR Research; 120Hz) and recorded for offline analyses at 1000 Hz. Visual stimuli were presented at a viewing distance of 51 cm on a Cambridge Research Systems display (1920 x 1080 pixels; 120 HZ refresh rate).

Monkeys were rewarded for detecting identity changes in visual objects that moved across the display while maintaining central fixation. Each trial began when the animal acquired and held a central fixation spot (0.25° radius; ° = degrees visual angle) on a uniform gray background. After 200 ms of fixation, a task-irrelevant static opaque occluder (4-8° diameter) appeared in the periphery. A single object stimulus (2-4° diameter) then appeared at a randomly selected start location positioned 15° from the occluder. After remaining stationary for 200 ms, the object moved towards the occluder at 15°/s for ∼1s. The object was initially fully visible (∼670 ms), then partially occluded (∼250 ms), and then fully occluded behind the occluder.

Task structure differed between animals. In one version (monkey Hd), the object stopped behind the occluder and remained fully occluded for ∼1 s; the occluder was then removed to reveal the object. In the other version (monkey Hb), the object continued moving behind the occluder along the same trajectory for ∼700 ms and then re-emerged on the opposite side.

On a subset of occluded trials (20-40%), the object’s identity was changed during the fully occluded interval. Animals were rewarded for shifting their gaze to the newly revealed object. On the remaining occluded trials, object identity was unchanged, and animals were rewarded for maintaining fixation. On an additional 20-30% of trials, no occluder was presented, and the object remained visible throughout, with otherwise matched timing.

In Task 1, four objects moved along four trajectories that intersected the occluder. The occluder contained gray textured line elements (line length 1-2°; 100-160 elements) with one of four orientations in sessions 1-4, and no line texture in sessions 5-7. In Task 2, six objects moved along two trajectories that intersected the occluder (clockwise or anticlockwise, co-eccentric), and a single occluder texture orientation was used throughout.

Trials were classified separately for occluded and visible conditions as hits, misses, false alarms, or correct rejections. A hit was defined as a gaze shift to the target on change trials; failure to respond on change trials was scored as a miss. A false alarm was defined as a gaze shift on no-change trials; maintaining fixation on no-change trials was scored as a correct rejection. Behavioral sensitivity was quantified using signal detection theory as

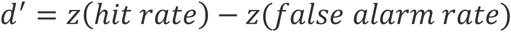

Where *z(.)* denotes the inverse cumulative distribution function of the standard normal distribution (MATLAB “norminv”).

### Surgical procedures and Electrophysiological Recordings

In each animal, a recording chamber was implanted over the superior temporal sulcus (STS) in either the left hemisphere (monkey Hd) or the right hemisphere (monkey Hb), centered ∼17 mm lateral to the midline and ∼7 mm posterior to ear-bar zero. A craniotomy was performed within the chamber to provide access to multiple visual areas along STS. All surgical procedures were performed using aseptic technique under general anesthesia, and postoperative analgesics were administered during recovery.

We recorded spiking activity from area MT, MST, FST using primate Neuropixels probes (IMEC). In each recording session, the dura was perforated with a 21-gauge pointed cannula mounted on an independent micromanipulator. A single Neuropixels NHP probe was then advanced through the cannula using either a hydraulic Narishge microdrive (monkey Hd) or a motorized drive systems with custom 3D-printed grid inserts (NAN instruments; monkey Hb). Recording sites were configured as 384 simultaneously active channels in a contiguous block (3.84mm span), providing dense sampling along the probe shank.

Probe trajectories spanned multiple cortical areas, as inferred from session-specific physiological signatures, including systematic progressions in receptive-field location across recording depth. After positioning, recordings were allowed to settle for ∼30 min before task onset to reduce drift.

### Data acquisition and spike sorting

Neuropixels probes digitized signals at the headstage and streamed separate bands for action potentials (AP; 300 Hz high-pass; 30 kHz sampling) and local field potential (LFP; 1 kHz low-pass; 2.5 kHz sampling). Neural activity was monitored online and saved to disk using SpikeGLX (https://billkarsh.github.io/SpikeGLX/).

Spike sorting was performed with Kilosort 4 (Pachitariu et al., 2024) to estimate spike times and assign spikes to templates (putative single units). Default Kilosort 4 parameters were used except for settings adjusted for Neuropixels NHP 1.0 probe and our recording environment: Ops.n_chan_bin=385; Ops.batch_size=180000; Ops.dminx=90; Ops.position_limit=500; Ops.max_channel_distance=160; Ops.nblocks=5. Sorted output was then manually curated in Phy (https://github.com/cortex-lab/phy) to remove units with very few spike counts or atypical waveforms and to perform minimal template merging/splitting. To remove potential double-counted spikes arising from residual fitting, we identified template pairs separated by < 50μm (∼5 channels) with spikes occurring within 5 samples (0.167ms), following established criteria (Siegle et al., 2021). For such double-counted events, spikes assigned to the smaller-amplitude template were excluded from subsequent analyses.

Furthermore, only neurons with a mean firing rate > 1 Hz during the late-occlusion epoch of occluded trials were included in subsequent analysis.

### Receptive field mapping

We mapped visual receptive fields (RF) by presenting a drifting Gabor probe at randomly selected locations on a 21(vertical) × 17 (horizontal) grid spanning 40° × 32°. The probe stimulus was a drifting Gabor patch (2° diameter; spatial frequency 0.5 cycle/°; drift speed 6°/ s; 100% Michelson contrast) presented for 100 ms with a 100 ms inter-stimulus interval. Each fixation trial contained 8-10 probes. Monkeys were rewarded with juice for maintaining fixation on a central spot (0.25° radius) throughout the trial.

For each neuron and each grid location, responses were quantified as the mean spike count from 50-250 ms after probe onset. This yielded a 21 × 17 response matrix per neuron. The matrix was then interpolated to 200 × 200 (MATLAB “interp2”) and smoothed with a 2D Gaussian filter (MATLAB “imgaussfilt”, *σ*=8). To compute a population RF map, each neuron’s response matrix was normalized by its maximum value and then averaged across neurons; the resulting map was visualized as a heat map (Figure 2b).

### Single-neuron and population firing rates

For each recording session and for each trajectory × object condition, we computed peri-stimulus time histograms (PSTHs) for all isolated neurons. Spike trains from 500 ms before to 3500 ms after fixation onset were segmented and binned at 1 ms, then averaged across all trials of the corresponding condition. The resulting PSTHs were convolved with a Gaussian kernel (σ = 150 ms) to obtain smoothed firing-rate time courses.

To enable comparison across neurons, each neuron’s smoothed responses were normalized by its session-wise maximum taken over all conditions and time points, yielding values in [0, 1]. For visualization, the normalized responses were displayed as heat maps for each trajectory × object condition, with neurons stacked along the y-axis in order of recorded cortical depth and time along the x-axis. For display only, the time axis was referenced to the onset of full occlusion (i.e., time zero corresponds to full occlusion onset; Figure 2b).

For each trajectory × object condition, we then computed a session-level population response by averaging the normalized responses across all neurons recorded in that session, resulting in a single population firing-rate trace per condition (Figure 2c).

### Classification of object identity and trajectory

For each recording session, we quantified population-level information about object identity and movement trajectory using linear decoding of simultaneously recorded STS neurons. Decoding was performed separately for visible and occluded conditions, using only behaviorally correct trials. To track the time course of encoded information, classifiers were trained at successive time points sampled every 25 ms, using spike counts computed within a 150 ms sliding window centered on each time point.

### Removal of task confounds

When decoding object identities during occlusion, movement trajectories (Task 1: four trajectories; Task 2: two trajectories) and, when applicable, occluder line orientations (Task 1 sessions 1-4: four orientations) were treated as nuisance variables. We constructed a trial-wise design matrix with one-hot encodings for trajectory and occluder orientation, as well as their interaction terms. In sessions without an occluder (Task 1 sessions 5-6), in Task 2 sessions with a single occluder, and in all visible conditions, occluder-related regressors were omitted. To control for slow session drift, we additionally included a trial-index time basis as regressors. For each time bin, we fit a ridge regression model (*α* = 3) from the design matrix to neural activity; the model-predicted component was subtracted, and the residual response were used for decoding.

When decoding trajectory, the same procedure was applied, except that object identity was treated as the nuisance variable.

### Decoders and cross-validation

We used multinomial logistic regression (scikit-learn “LogisticRegression”, solver= “lbfgs”, max_iter=2000) with L2 regularization (C=1) to classify object identity (Task 1: four classes; Task 2: six classes) or trajectory. Performance was evaluated with 5-fold cross-validation and reported as percentage correct on held-out trials. Any preprocessing steps (including the confound-regression procedure described above) were fit using training folds only and then applied to both training and testing folds to prevent information leakage.

Because the number of classes differed across tasks/sessions, decoding performance was additionally expressed as a chance-corrected relative accuracy:

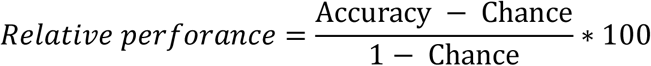

where Chance is the theoretical chance level for the corresponding number of classes.

### Decay timescale estimation

We modeled the time course of decoding performance with a delayed exponential decay:

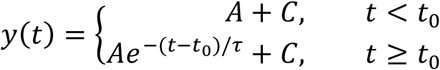

where *t_0_* is the decay onset, *τ* is the post-onset decay time constant, and *C* is the asymptotic value. Parameters were estimated by nonlinear least-squares regression (MATLAB “nlinfit”). Confidence intervals for fitted parameters were obtained from the estimated parameter covariance matrix returned by “nlinfit” using “nlparci” (95% CI). To summarize the overall decay timescale, we additionally report *t_95_* defined as the time from full occlusion onset to 95% completion of the decay toward the asymptote: *t_95_= t_0_* + *τ \** ln (1/0.05) (i.e., the time at which the fitted response is within 5% of its total drop from *A*+*C* to *C*).

### Statistical significance for decoding

Significance was assessed with a permutation test in which class labels were randomly shuffled within each trajectory × occluder condition (to preserve any confounding structure). For each time bin, we performed 100 shuffles and computed a p value as the fraction of shuffled accuracies exceeded the unshuffled accuracy.

### Late-occlusion window analysis

To increase statistical power, we repeated the decoding analysis using, for each neuron, the mean firing rate within a predefined late-occlusion window for occluded trials and the corresponding time window for visible trials. The same late-occlusion window definition was used for all subsequent CCG analyses.

For occluded trials, the late-occlusion window was defined relative to full-occlusion onset as 0.57-1.07s (500 ms duration) in Task 1 and 0.47-0.77s (300 ms duration) in Task 2, chosen to capture the period after the initial pre-occlusion response had decayed to an apparent asymptote. For visible trials, the corresponding window was shorter because object-identity changes (when present) became visible earlier: chosen as 0.57-0.87s (300 ms duration) for Task 1 and 0.47-0.67s (200 ms) for Task 2, referenced to the time of full occlusion onset in the matched occluded-trial timeline.

### Cross-correlograms (CCGs) analysis

To test whether identity information for hidden objects was expressed in coordinated spiking at fine temporal scales (<100ms), we computed cross-correlograms (CCGs) for all pairs of simultaneously recorded neurons, using spikes from the late-occlusion epoch. For each neuronal pair *j* (reference) and *k* (target), we computed the CCG at time lag τ as:

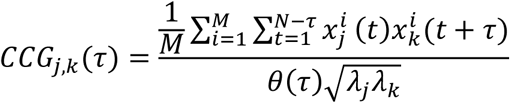

where 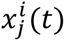 and 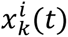 are binned spike trains (1ms bins; value 1 if a spike occurred in bin *t*, otherwise 0) for trial *i*; *M* is the number of trials and *N* is the number of bins in the analysis window. The factor *θ*(*τ*) = *N* − |*τ*| is a triangular correction for the decreasing temporal overlap at larger |*τ*|. *λ*_*j*_ and *λ*_*k*_ are the mean firing rates of neuron *j* and *k*, computed over the same bins used for the CCG. Normalization by 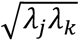 reduces firing-rate-dependent effects on CCG amplitude (Bair et al., 2001; Kohn & Smith, 2005). To remove correlations attributable to stimulus locking and slower shared fluctuations, we computed a jitter-corrected CCG:

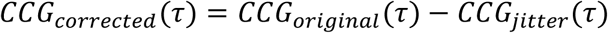

*CCG_jitter_* is the expected CCG under spike-time jittering within a fixed window (Harrison & Geman, 2009; Smith & Kohn, 2008). Jittering preserves both the PSTH across trials and spike counts within each jitter window within each trial, thereby removing correlations on timescales longer than the jitter window while retaining fast interactions. We used a 25 ms jitter window following prior studies (Jia et al., 2013; Siegle et al., 2021).

### Identification of significant CCGs and CCG-derived metrics

We classified a neuronal pair as exhibiting significant fast-timescale interaction if the peak of CCG_corrected_ occurred within ±10 ms of zero lag and exceeded 7 standard deviations above a noise distribution (Siegle et al., 2021). The noise distribution was defined from the flanks of the jitter-corrected CCG: {*CCG*_*corrected*_ (*τ*)|50 ≤ |*τ*| ≤ 100 *ms*}.

From each CCG we extracted two summary measures. One is CCG peak amplitude, calculated as the maximum value of *CCG*_*corrected*_ (*τ*) within ±10 ms, after subtracting the mean of the noise distribution (yielding a baseline-corrected “excess coincidence” measure). The second measure is the peak lag, or the τ at which the peak occurred. Peak lag was defined as positive when reference neuron *j* led target neuron *k*, negative when *j* lagged *k*, and zero for synchronous firing.

In analyses that included **all** neuronal pairs (i.e., significant and non-significant; Figure 4c, Figure 5c), we computed the peak amplitude and peak lag identically, except that we did not require the significance criterion; for non-significant pairs, the peak was simply the maximum within ±10 ms (again baseline-corrected by the noise distribution); and the peak lag was τ at which the peak occurred within ±10 ms of zero lag.

### Trial matching for comparing object-specific differences in CCG structure

Within each recording session, we compared CCGs between object identities by considering all pairwise object comparisons (Task 1: 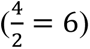 comparisons; Task 2: 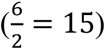 comparisons). For each object comparison, we controlled for differences in trial counts across trajectory × occluder conditions by matching trial numbers between the two objects within each trajectory × occluder cell. Specifically, within each cell we subsampled (without replacement) the smaller of the two trial counts, then pooled selected trials across trajectory × occluder cells. This ensured that, for every trajectory × occluder condition, the two objects contributed equal number of trials, preventing confounding by object-dependent trial imbalances. For each object comparison, we repeated the subsampling procedure 20 times and averaged the resulting estimates, yielding one CCG per object for each neuronal pair.

### Population distance using significant CCGs

Because significance could differ between the two objects for a given neuronal pair, for each object comparison, we first defined an analysis set consisting of neuronal pairs with a significant CCG for at least one of two objects. For this set, we formed a vector of baseline-corrected peak amplitude across neuronal pairs for each object, and computed the Euclidean distance between the two peak-amplitude vectors, yielding a session-level population distance for that object comparison. To assess significance, we generated a null distribution by shuffling object labels within each of the 20 subsamples (50 shuffles per subsample; 1000 shuffles total). For each shuffle, we recomputed the CCGs and population-distance metric using the same set of neuronal pairs as in the unshuffled analysis. We quantified the observed distance as a z-score relative to the shuffle distribution, and evaluated significance using a permutation p-value, defined as the fraction of shuffled distances greater than (i.e., exceeding) the observed distance (Figure 4b).

### Population shift using all CCGs

To assess object-dependent shifts without restricting to significant CCGs, for each object comparison within each recording session, we included all neuronal pairs considered in the significant-CCG analysis but did not apply the significance criterion. All other procedures were identical: trial counts were matched between objects within each trajectory × occluder cell as described above; CCGs were computed with the same binning, lag window, jitter correction, and baseline estimation; and subsampling was repeated 20 times with estimates averaged to yield one CCG per object per neuronal pair.

For each neuronal pair, we extracted the baseline-corrected near-zero peak amplitude of the jitter-corrected CCG (maximum within ±10 ms after subtracting the mean of the noise flanks) for both objects in the comparison, and formed an object-comparison difference Δ = peak(object A) - peak(object B). Under the null hypothesis of no object-specific structure, the distribution of Δ across neuronal pairs should be centered at zero. For each object comparison, we assessed a population shift with a Wilcoxon signed-rank test on the set of Δ values. As a scale-free summary effect size, we reported the signed standardized mean of the differences (Cohen’s dz = mean(Δ)/SD (Δ)). To pool across object comparisons and sessions, we used the absolute value of dz (Figure 5c).

## Acknowledgements

The authors would like to thank Bob Schneeveis for machining assistance, Donatas Jonikaitis for developing behavioral control system, Ethan B. Trepka, Matthew F. Panichello, Sharif Saleki, Ruobing Xia, Shreyas Muralidharan, Francesco Lanfranchi for helpful comments. This work was supported by NIH grants EY014924 and NS11662302, and a Ben Barres Professorship to T.M.

## References

Assad, J. A., & Maunsell, J. H. (1995). Neuronal correlates of inferred motion in primate posterior parietal cortex. Nature, 373(6514), 518–521. 10.1038/373518a0

Bair, W., Zohary, E., & Newsome, W. T. (2001). Correlated firing in macaque visual area MT: time scales and relationship to behavior. J Neurosci, 21(5), 1676–1697. 10.1523/JNEUROSCI.21-05-01676.2001

Baker, C. I., Keysers, C., Jellema, T., Wicker, B., & Perrett, D. I. (2001). Neuronal representation of disappearing and hidden objects in temporal cortex of the macaque. Exp Brain Res, 140(3), 375–381. 10.1007/s002210100828

Bichot, N. P., Heard, M. T., DeGennaro, E. M., & Desimone, R. (2015). A Source for Feature-Based Attention in the Prefrontal Cortex. Neuron, 88(4), 832–844. 10.1016/j.neuron.2015.10.001

Bichot, N. P., Xu, R., Ghadooshahy, A., Williams, M. L., & Desimone, R. (2019). The role of prefrontal cortex in the control of feature attention in area V4. Nat Commun, 10(1), 5727. 10.1038/s41467-019-13761-7

Bisley, J. W., Zaksas, D., Droll, J. A., & Pasternak, T. (2004). Activity of neurons in cortical area MT during a memory for motion task. J Neurophysiol, 91(1), 286–300. 10.1152/jn.00870.2003

Born, R. T., & Bradley, D. C. (2005). Structure and function of visual area MT. Annu Rev Neurosci, 28, 157–189. 10.1146/annurev.neuro.26.041002.131052

Christophel, T. B., Klink, P. C., Spitzer, B., Roelfsema, P. R., & Haynes, J. D. (2017). The Distributed Nature of Working Memory. Trends Cogn Sci, 21(2), 111–124. 10.1016/j.tics.2016.12.007

Cornelissen, F. W., Peters, E. M., & Palmer, J. (2002). The Eyelink Toolbox: eye tracking with MATLAB and the Psychophysics Toolbox. Behav Res Methods Instrum Comput, 34(4), 613–617. 10.3758/bf03195489

de Bois, S. T., & Novak, M. A. (1994). Object permanence in rhesus monkeys (Macaca mulatta). Journal of Comparative Psychology, 108(4), 318–327. 10.1037/0735-7036.108.4.318

Dotson, N. M., Hoffman, S. J., Goodell, B., & Gray, C. M. (2018). Feature-Based Visual Short-Term Memory Is Widely Distributed and Hierarchically Organized. Neuron, 99(1), 215–226 e214. 10.1016/j.neuron.2018.05.026

Fuster, J. M., & Alexander, G. E. (1971). Neuron activity related to short-term memory. Science, 173(3997), 652–654. 10.1126/science.173.3997.652

Harrison, M. T., & Geman, S. (2009). A rate and history-preserving resampling algorithm for neural spike trains. Neural Comput, 21(5), 1244–1258. 10.1162/neco.2008.03-08-730

Harrison, S. A., & Tong, F. (2009). Decoding reveals the contents of visual working memory in early visual areas. Nature, 458(7238), 632–635. 10.1038/nature07832

Huang, J., Wang, T., Dai, W., Li, Y., Yang, Y., Zhang, Y.,…Xing, D. (2024). Neuronal representation of visual working memory content in the primate primary visual cortex. Sci Adv, 10(24), eadk3953. 10.1126/sciadv.adk3953

Jia, X., Tanabe, S., & Kohn, A. (2013). gamma and the coordination of spiking activity in early visual cortex. Neuron, 77(4), 762–774. 10.1016/j.neuron.2012.12.036

Jonikaitis, D., Xia, R., & Moore, T. (2025). Robust encoding of stimulus-response mapping by neurons in visual cortex. Proc Natl Acad Sci U S A, 122(9), e2408079122. 10.1073/pnas.2408079122

Kleiner, M., Brainard, D., & Pelli, D. (2007). What’s new in Psychtoolbox-3? Perception, 36, 14–14.

Kohn, A., & Smith, M. A. (2005). Stimulus dependence of neuronal correlation in primary visual cortex of the macaque. J Neurosci, 25(14), 3661–3673. 10.1523/JNEUROSCI.5106-04.2005

Kravitz, D. J., Saleem, K. S., Baker, C. I., & Mishkin, M. (2011). A new neural framework for visuospatial processing. Nat Rev Neurosci, 12(4), 217–230. 10.1038/nrn3008

Lundqvist, M., Herman, P., & Miller, E. K. (2018). Working Memory: Delay Activity, Yes! Persistent Activity? Maybe Not. J Neurosci, 38(32), 7013–7019. 10.1523/JNEUROSCI.2485-17.2018

Mendoza-Halliday, D., Torres, S., & Martinez-Trujillo, J. C. (2014). Sharp emergence of feature-selective sustained activity along the dorsal visual pathway. Nat Neurosci, 17(9), 1255–1262. 10.1038/nn.3785

Miller, E. K., Erickson, C. A., & Desimone, R. (1996). Neural mechanisms of visual working memory in prefrontal cortex of the macaque. J Neurosci, 16(16), 5154–5167. 10.1523/JNEUROSCI.16-16-05154.1996

Miller, E. K., Li, L., & Desimone, R. (1993). Activity of neurons in anterior inferior temporal cortex during a short-term memory task. J Neurosci, 13(4), 1460–1478. 10.1523/JNEUROSCI.13-04-01460.1993

Miyashita, Y., & Chang, H. S. (1988). Neuronal correlate of pictorial short-term memory in the primate temporal cortex. Nature, 331(6151), 68–70. 10.1038/331068a0

Mongillo, G., Barak, O., & Tsodyks, M. (2008). Synaptic theory of working memory. Science, 319(5869), 1543–1546. 10.1126/science.1150769

Motter, B. C. (1994). Neural correlates of feature selective memory and pop-out in extrastriate area V4. J Neurosci, 14(4), 2190–2199. 10.1523/JNEUROSCI.14-04-02190.1994

O’Herron, P. J., & von der Heydt, R. (2009). Short-term memory for figure-ground organization in the visual cortex. Neuron, 61(5), 801–809. (Not in File)

Pachitariu, M., Sridhar, S., Pennington, J., & Stringer, C. (2024). Spike sorting with Kilosort4. Nat Methods, 21(5), 914–921. 10.1038/s41592-024-02232-7

Panichello, M. F., & Buschman, T. J. (2021). Shared mechanisms underlie the control of working memory and attention. Nature, 592(7855), 601–605. 10.1038/s41586-021-03390-w

Panichello, M. F., Jonikaitis, D., Oh, Y. J., Zhu, S., Trepka, E. B., & Moore, T. (2024). Intermittent rate coding and cue-specific ensembles support working memory. Nature, 636(8042), 422–429. 10.1038/s41586-024-08139-9

Pearson, J. (2019). The human imagination: the cognitive neuroscience of visual mental imagery. Nat Rev Neurosci, 20(10), 624–634. 10.1038/s41583-019-0202-9

Piaget, J. (1954). The development of object concept. In M. Cook (Ed.), The construction of reality in the child. (pp. 3–96). Basic Books/Hachette Book Group. 10.1037/11168-001

Sakai, K., & Miyashita, Y. (1991). Neural organization for the long-term memory of paired associates. Nature, 354(6349), 152–155. 10.1038/354152a0

Schaffer, A., Widdig, A., Holland, R., & Amici, F. (2024). Evidence of object permanence, short-term spatial memory, causality, understanding of object properties and gravity across five different ungulate species. Sci Rep, 14(1), 13718. 10.1038/s41598-024-64396-8

Schlack, A., & Albright, T. D. (2007). Remembering visual motion: neural correlates of associative plasticity and motion recall in cortical area MT. Neuron, 53(6), 881–890. 10.1016/j.neuron.2007.02.028

Serences, J. T. (2016). Neural mechanisms of information storage in visual short-term memory. Vision Res, 128, 53–67. 10.1016/j.visres.2016.09.010

Serences, J. T., Ester, E. F., Vogel, E. K., & Awh, E. (2009). Stimulus-specific delay activity in human primary visual cortex. Psychol Sci, 20(2), 207–214. 10.1111/j.1467-9280.2009.02276.x

Siegle, J. H., Jia, X., Durand, S., Gale, S., Bennett, C., Graddis, N.,…Koch, C. (2021). Survey of spiking in the mouse visual system reveals functional hierarchy. Nature, 592(7852), 86–92. 10.1038/s41586-020-03171-x

Smith, M. A., & Kohn, A. (2008). Spatial and temporal scales of neuronal correlation in primary visual cortex. J Neurosci, 28(48), 12591–12603. 10.1523/JNEUROSCI.2929-08.2008

Stokes, M. G. (2015). ’Activity-silent’ working memory in prefrontal cortex: a dynamic coding framework. Trends Cogn Sci, 19(7), 394–405. 10.1016/j.tics.2015.05.004

Super, H., Spekreijse, H., & Lamme, V. A. (2001). A neural correlate of working memory in the monkey primary visual cortex. Science, 293(5527), 120–124. 10.1126/science.1060496

van Kerkoerle, T., Self, M. W., & Roelfsema, P. R. (2017). Layer-specificity in the effects of attention and working memory on activity in primary visual cortex. Nat Commun, 8, 13804. 10.1038/ncomms13804

Woloszyn, L., & Sheinberg, D. L. (2009). Neural dynamics in inferior temporal cortex during a visual working memory task. J Neurosci, 29(17), 5494–5507. 10.1523/JNEUROSCI.5785-08.2009

Yiling, Y., Klon-Lipok, J., Shapcott, K., Lazar, A., & Singer, W. (2024). Dynamic fading memory and expectancy effects in the monkey primary visual cortex. Proc Natl Acad Sci U S A, 121(8), e2314855121. 10.1073/pnas.2314855121

Zaksas, D., & Pasternak, T. (2006). Directional signals in the prefrontal cortex and in area MT during a working memory for visual motion task. J Neurosci, 26(45), 11726–11742. 10.1523/JNEUROSCI.3420-06.2006

